# β-amyloid−driven synaptic depression requires PDZ protein interaction at AMPA-receptor subunit GluA3

**DOI:** 10.1101/2021.10.03.462970

**Authors:** Niels R. Reinders, Sophie van der Spek, Remco V. Klaassen, Karin J. Koymans, Ka Wan Li, August B. Smit, Helmut W. Kessels

## Abstract

Soluble oligomeric amyloid-β (Aβ) is a prime suspect to cause cognitive deficits in Alzheimer’s disease and weakens synapses by removing AMPA-type glutamate receptors (AMPARs). We show that synapses of CA1 pyramidal neurons become vulnerable to Aβ when they express AMPAR subunit GluA3. We found that Aβ-oligomers reduce the levels of GluA3 immobilized at spines, indicating they deplete GluA3-containing AMPARs from synapses. These Aβ-driven effects critically depended on the PDZ-binding motif of GluA3. When GluA3 was expressed with a single amino acid mutation in its PDZ-binding motif that prevents GRIP binding, it did not end up at spines and Aβ failed to trigger synaptic depression. GluA3 with a different point mutation in the PDZ-motif that leaves GRIP-binding intact but prevents its endocytosis, was present at spines in normal amounts but was fully resistant to effects of Aβ. Our data indicate that Aβ-mediated synaptic depression requires the removal of GluA3 from synapses. We propose that GRIP-detachment from GluA3 is a critical early step in the cascade of events through which Aβ accumulation causes a loss of synapse.

## Introduction

Synapse loss is considered a mayor cause for cognitive deficits in Alzheimer’s disease (AD), and is strongly correlated with AD symptoms (de Wilde et al., 2016, Selkoe, 2002). Studies in rodent models have shown that the accumulation of amyloid-β (Aβ) leads to a loss of synapses as a consequence of the removal of AMPA-receptors (AMPARs) from synapses (Jurado, 2018, Guntupalli, Widagdo & Anggono, 2016). The majority of AMPARs in CA1 pyramidal neurons consist of either subunits GluA1 and GluA2 (GluA1/2) or GluA2 and GluA3 (GluA2/3) (Wenthold et al., 1996). The carboxy-terminal tail (c-tail) of GluA2 and GluA3 are similar in sequence and share an identical PDZ-binding motif. Through this motif, GluA2 and GluA3 interact with PDZ-containing proteins GRIP and PICK1. GRIP-binding controls the transport of AMPARs into dendrites and their stabilization at synapses (Setou et al., 2002, Osten et al., 2000), whereas interaction with PICK1 promotes AMPAR endocytosis and their degradation in lysosomes (Koszegi, Fiuza & Hanley, 2017, Kim et al., 2001, Perez et al., 2001, Fiuza et al., 2017). The PDZ-binding motif can be phosphorylated by PKCα, which disrupts its interaction with GRIP but not with PICK1. As such, PKCα can regulate GRIP/PICK1-mediated cycling of AMPARs in and out of synapses (Moretto, Passafaro, 2018, Hanley, 2018). Notably, PKCα phosphorylation of AMPARs and their endocytosis through interaction with PICK1 are involved in Aβ-driven synaptic depression and spine loss (Hsieh et al., 2006, Alfonso, S. et al., 2014, Alfonso, S. I. et al., 2016). Because PICK1 can interact with GluA2 and GluA3, both GluA1/2s and GluA2/3s should theoretically be susceptible to Aβ-mediated removal from synapses. However, we previously found that neurons are fully resistant to Aβ-mediated synaptic depression when they lack GluA3 (Reinders et al., 2016), implying that GluA3-containing AMPARs are selectively targeted by Aβ. We therefore set out to investigate how GluA3 influences the vulnerability of neurons for Aβ-driven synaptic depression.

## Results and Discussion

### GluA3 expression sensitizes CA1 neurons to Aβ-mediated synaptic depression

We expressed GFP-GluA3 in a subset of CA1 neurons within organotypic hippocampal slices isolated from GluA3-knockout mice by Sindbis viral transfection, which did not affect the health of neurons (Fig. S1) (Uyaniker et al., 2019). AMPAR subunits expressed in CA1 neurons by viral transduction accumulate in cell bodies, but appear in dendrites at near-physiological levels (Kessels et al., 2009). Similarly, 48 hours after GFP-GluA3 transfection, levels were highest in cell bodies and lower in dendrites (Fig. S2). GFP-GluA3 was detectable at the majority of spines (Fig. 1A,B). AMPARs have been shown to freely diffuse across the extra-synaptic spine surface and stay largely immobilized at synapses (Triller, Choquet, 2005, Ehlers et al., 2007). To assay AMPAR mobility on spines, GluA3 was tagged with a pH-sensitive GFP variant (SEP) to selectively visualize surface receptors (Kopec et al., 2006) and we measured fluorescence recovery after photobleaching (FRAP) at individual spines (Fig. S3A) (Makino, Malinow, 2009). The level of FRAP stabilized at ∼45% of the initial spine fluorescence (Fig. 1C), indicating that on average 55% of recombinant GluA3 at the spine surface was immobilized. GFP-GluA3 expression did not alter synaptic currents measured by recording either miniature excitatory post-synaptic currents (mEPSCs) (Fig. 1D) or currents evoked by stimulation of Schaffer collateral inputs (eEPSC) (Fig. 1E).

**Figure 1.**
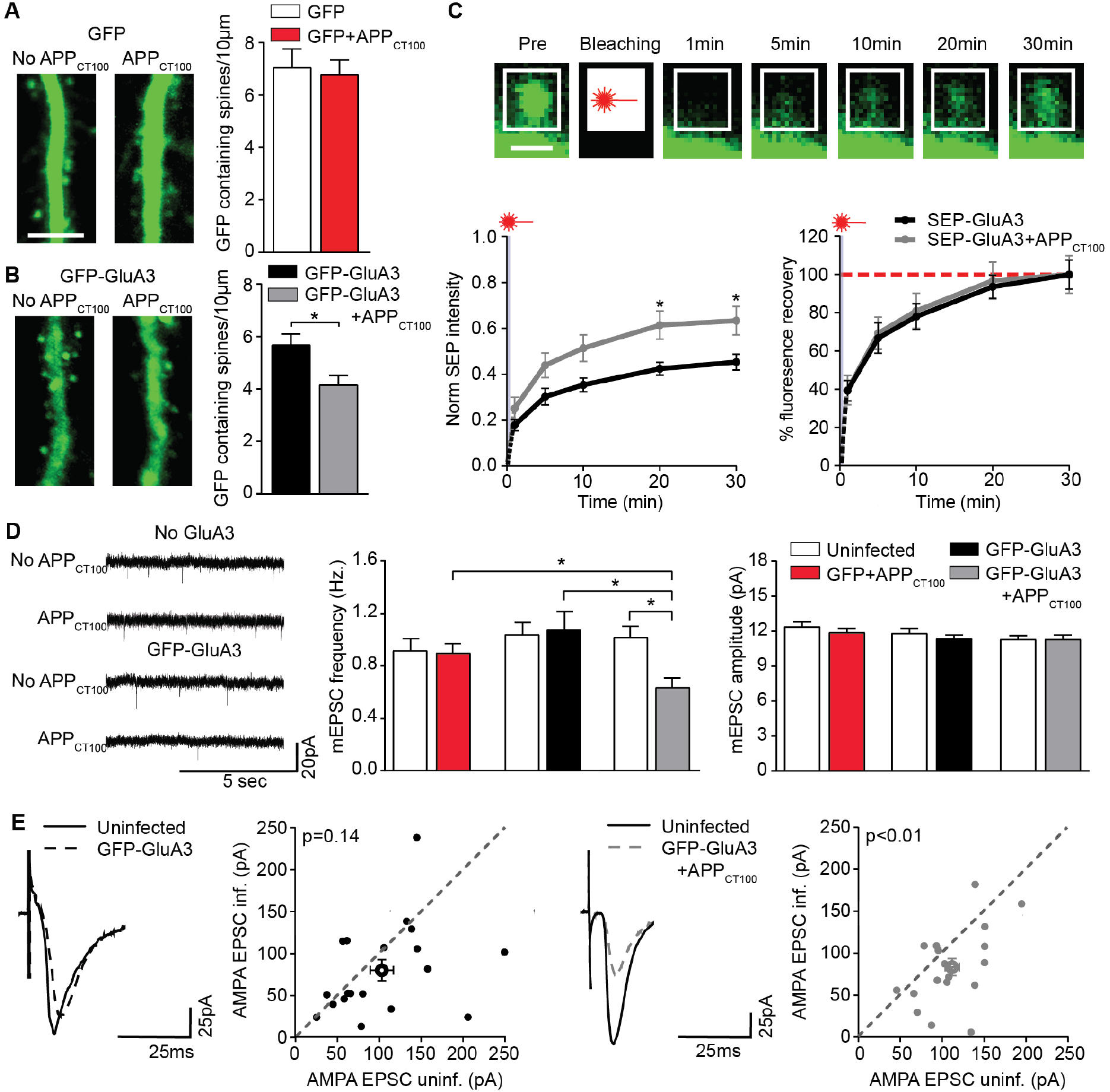
Neuronal expression of GluA3 is sufficient for Aβ to impair synaptic function. (A) Spine density in GluA3-KO neurons was unaffected by GFP+APP_CT100_ expression (p=0.778; GFP n=19, GFP+APP_CT100_ n=15; example images scale bar: 5 μm). (B) GluA3-KO neurons expressing GFP-GluA3 showed a lower density of GFP-containing spines when APP_CT100_ was co-expressed (p= 0.008; GFP-GluA3 n=25, GFP-GluA3+APP_CT100_ n=29). (C top) Time series of SEP-GluA3 expressing dendritic spine before and after fluorescence bleaching (scale bar: 1 µm). (left) FRAP of dendritic spines expressing SEP-GluA3 (black, n=22) was increased by APP_CT100_ co-expression, demonstrating a reduced immobile fraction of SEP-GluA3 (t20 and 30 p=0.015; grey, n=22). (right) FRAP speed of SEP-GluA3 was unaffected by APP_CT100_ co-expression (t5, 10 and 20 p=0.989). (D) Example mEPSC traces of GluA3-KO neurons with or without APP_CT100_ and/or GFP-GluA3 expression. (center) Only the combined expression of GFP-GluA3 with APP_CT100_ lowered mEPSC frequency (ANOVA p=0.002: GFP-GluA3+CT100 vs GFP+APP_CT100_ p=0.016; GFP-GluA3 vs GFP-GluA3+CT100 p=0.008; uninf. vs GFP-GluA3+CT100 p= 0.002; uninf. conditions were not compared) but not (right) mEPSC amplitude (ANOVA p=0.433; uninf. n=24, GFP+APP_CT100_ n=29; uninf. n=29, GFP-GluA3 n=30; uninf. n= 29, GFP-GluA3+APP_CT100_ n= 27). (E) Example traces and dot plots (filled dots represent individual dual recording, open dots denote averages) of simultaneous dual EPSC recordings from (left) neighboring GFP-GluA3 infected and uninfected GluA3-KO neurons showed no significant synaptic depression (paired t-test; n=18) (right) unless APP_CT100_ was co-expressed (paired t-test; n=19). Data are mean ± SEM. *p < 0.01.

We then studied the role of GluA3 in Aβ-mediated synaptic depression by expressing APP_CT100_, the β-secretase product of APP. APP_CT100_-expression leads to synaptic depression and spine loss as a consequence of oligomeric Aβ production in wild-type CA1 neurons (Kessels, Nabavi & Malinow, 2013), but fails to do so in GluA3-deficient neurons (Reinders et al., 2016). Upon co-expression of GFP-GluA3 with APP_CT100_ in GluA3-deficient neurons, the number of GFP-containing spines decreased (Fig. 1B). FRAP analysis showed that co-expression of APP_CT100_ lowered the fraction of immobilized GluA3 at spines by 33% without affecting the velocity of mobile GluA3 on the spine surface (Fig. 1C). Co-expression of APP_CT100_ with GFP-GluA3 caused a significant decrease in mEPSC frequency (Fig. 1D) and eEPSCs amplitude (Fig. 1E). These data indicate that Aβ requires the presence of GluA3 at post-synaptic neurons to trigger synaptic depression.

### Aβ-mediated synaptic depression requires GRIP binding to GluA3

We next investigated the role of PDZ-protein interactions with GluA3 in the effects of Aβ on synapses. Within the PDZ-binding motif of GluA3, we mutated serine 885 to alanine (GluA3_S885A_), which abolished interaction with GRIP (Fig. S4). In comparison to GFP-GluA3, GFP-GluA3_S885A_ fluorescence levels were substantially lower at apical dendrites relatively to cell bodies (Fig. S2), and was detected at only a small proportion of spines (Fig 2A), prohibiting reliable FRAP analysis. GFP-GluA3_S885A_ expression also did not alter synaptic AMPAR currents (Fig. 2B,C). These data are in line with the notion that AMPARs need to bind GRIP to be stably expressed at synapses (Osten et al., 2000, Setou et al., 2002). To test whether GluA3 requires binding to GRIP for Aβ to cause synaptic depression, we co-expressed GFP-GluA3_S885A_ with APP_CT100_ in GluA3-deficient neurons. Regardless of APP_CT100_ co-expression, GFP containing spines were equally scarce (Fig. 2A) and synaptic currents remained unaffected, as reflected by average mEPSC frequency (Fig. 2B) or eEPSC amplitude (Fig. 2C). This observation that expression of GFP-GluA3_S885A_ is insufficient for re-sensitizing synapses to APP_CT100_ suggests it is not the overexpression of GluA3 at cell bodies that renders synapses vulnerable to Aβ (e.g. by disrupting assembly of endogenous AMPAR subunits) but that GluA3 needs to be transported by GRIP to dendrites and synapses.

**Figure 2.**
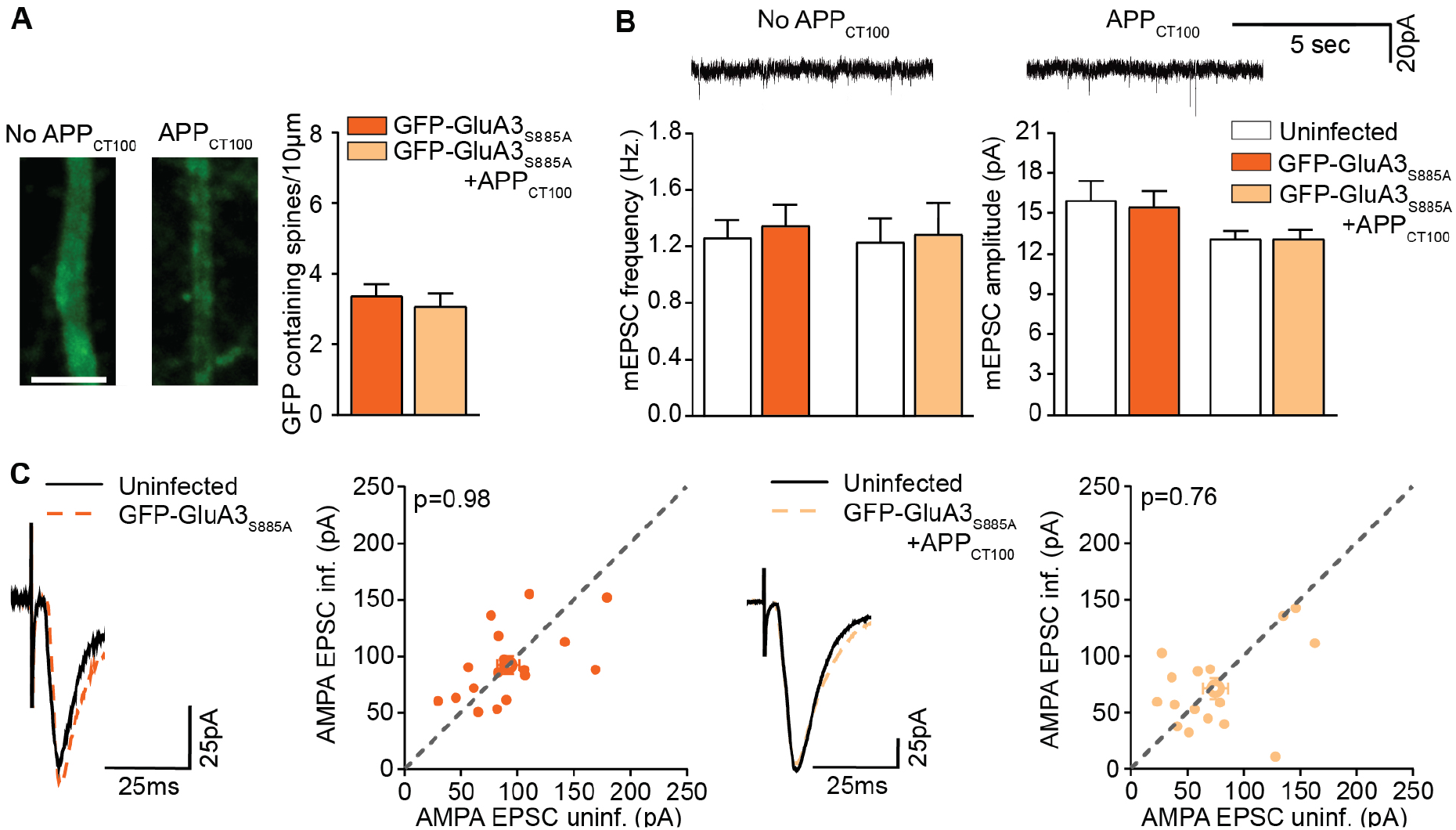
Neuronal expression of GluA3_S885A_ does not sensitize GluA3-KO neurons to Aβ. (A) GFP-GluA3_S885A_ expressing GluA3-KO dendrites showed a low density of GFP-containing spines, which was unchanged by APP_CT100_ co-expression (p=0.536; GFP-GluA3_S885A_ n=19, GFP-GluA3_S885A_+APP_CT100_ n=17; example images scale bar: 5 μm). (B top) Example mEPSC traces of GluA3-KO neurons expressing GFP-GluA3_S885A_ with or without APP_CT100_. (left) GFP-GluA3_S885A_ with or without APP_CT100_ did not affect mEPSC frequency (unpaired t-test vs. uninf.) or (right) amplitude (uninf. n=22, GFP-GluA3_S885A_ n=23; uninf. n=25, GFP-GluA3_S885A_ +APP_CT100_ n=29) (unpaired t-test vs. uninf.). (C) Example traces and dot plots (filled dots represent individual dual recording, open dots denote averages) of dual EPSC recordings from (left) GFP-GluA3_S885A_ expressing GluA3-KO neurons and their uninfected neighbor showed no synaptic depression (paired t-test; n=17), (right) similar to those co-expressing APP_CT100_ (paired t-test; n=16). Data are mean ± SEM.

### GluA3 PDZ-binding motif is required for Aβ-induced synaptic depression

We then generated a different mutation in the PDZ-binding motif of GluA3 by substituting lysine 887 with alanine (GluA3_K887A_), which disrupts the PKCα recognition site (Kreegipuu et al., 1998) while preserving its interaction with GRIP (Fig. S4). Upon expression of GFP-GluA3_K887A_ in GluA3-deficient neurons, GFP was detected in dendrites and at spines (Fig 3A and S1). FRAP analysis showed that 43% of GluA3_K887A_ was immobilized at spines (Fig 3B). Expression of GFP-GluA3_K887A_ led to a small decrease in mEPSC amplitude but not in mEPSC frequency (Fig 3C) and no change in eEPSC amplitude (Fig 3D). Directly comparing expression of GFP-GluA3 to GFP-GluA3_K887A_ indicates that this single amino acid mutation did not significantly alter the transport of GluA3 into spines (Fig. 4A), the fraction of GluA3 immobilized at spines (Fig. 4B) or synaptic currents (Fig. 4C-E). However, when we co-expressed GFP-GluA3_K887A_ with APP_CT100_, we observed no loss in GFP-containing spines (Fig. 3A), no change in the fraction of immobile GluA3_K887A_ (Fig. 3B), and no decrease in mEPSC frequency, amplitude (Fig. 3C) or eEPSC amplitude (Fig. 3D). Thus, expressing either GFP-GluA3_K887A_ or GFP-GluA3 made a significant difference when co-expressing APP_CT100_: a single amino acid in the PDZ-binding motif of GluA3 determined whether Aβ-oligomers were capable of decreasing the fraction of GluA3-containing AMPARs immobilized at spines and of reducing synaptic AMPAR currents (Fig. 4). These data suggest that Aβ-oligomers can only trigger synaptic depression when PDZ-protein interactions at GluA3-containing AMPARs are intact and can facilitate endocytosis.

**Figure 3.**
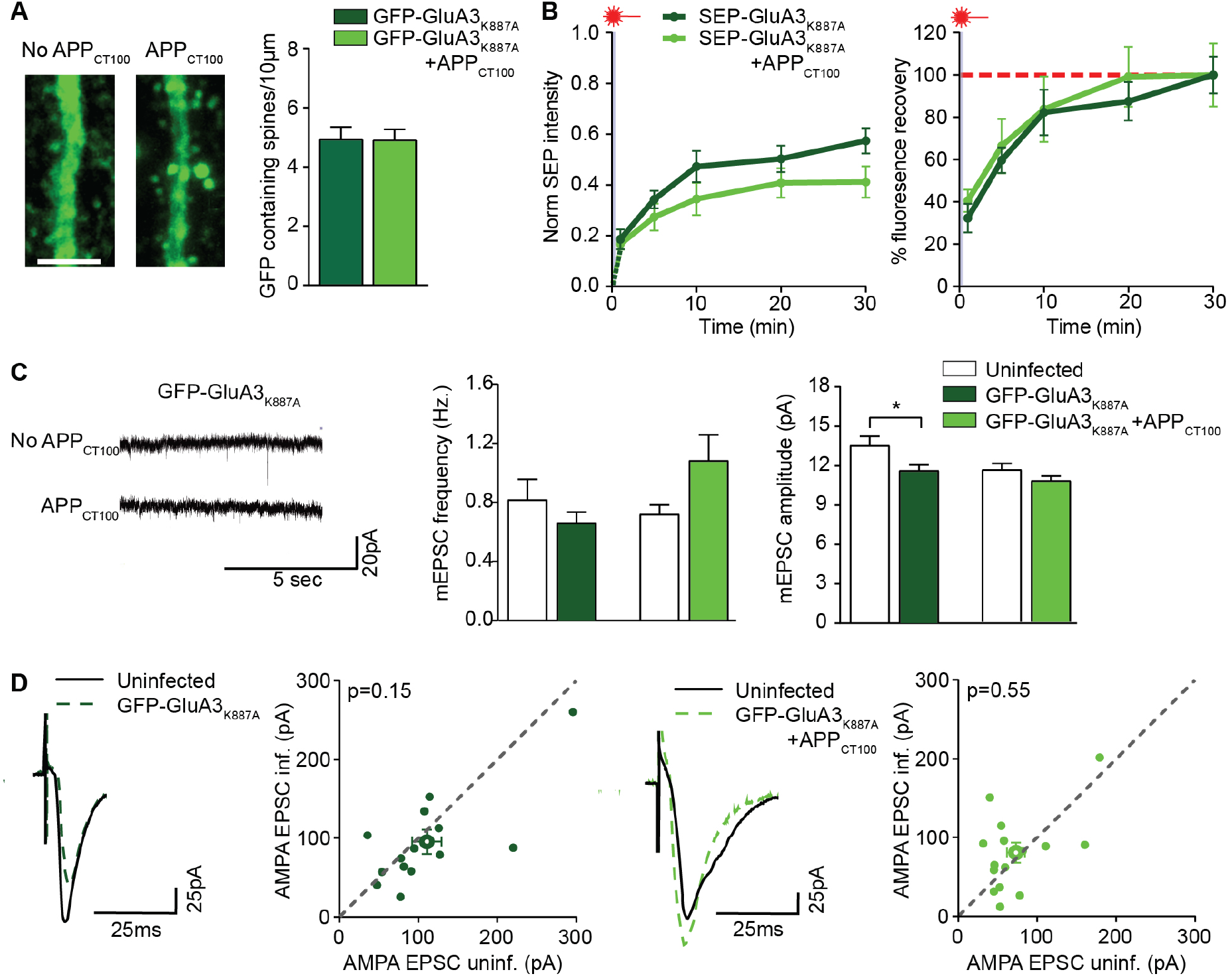
Neuronal expression of GluA3_K887A_ does not sensitize GluA3-KO neurons to Aβ. (A) Density of GFP-containing spines on GFP-GluA3_K887A_ expressing GluA3-KO dendrites was unchanged by APP_CT100_ co-expression (GFP-GluA3_K887A_ n=31, GFP-GluA3_K887A_ +APP_CT100_ n=27; example images scale bar: 5 μm). (B) In dendritic GluA3-KO spines expressing SEP-GluA3_K887A_ (dark green, n=12), the co-expression of APP_CT100_ co-expression (light green, n=13) did not affect FRAP (left; t20 p=0.241; t30 p=0.091) or its speed (right; t5 p=0.879; t10 p=0.936; t=20 p=0.879). (C left) Example mEPSC traces of GluA3-KO neurons expressing GFP-GluA3_K887A_ with or without APP_CT100_. (center) Expression of GFP-GluA3_K887A_ with (p=0.665) or without APP_CT100_ (p=0.59) did not affect mEPSC frequency (unpaired t-test vs. uninf.). (right) GFP-GluA3_K887A_ expression lowered mEPSC amplitude (p=0.0278) but not when APP_CT100_ was co-expressed (unpaired t-test vs. uninf. p=0.159; uninf n=23, GFP-GluA3_K887A_ n=28; uninfected n=27, GFP-GluA3_K887A_ + APP_CT100_ n=31). (D) Example traces and dot plots (filled dots represent individual paired recording, open dots denote averages) of (left) paired EPSC recordings from GFP-GluA3_K885A_ expressing GluA3-KO neurons and their uninfected neighbor showed no synaptic depression (paired t-test; n=15), (right) similar to those co-expressing APP_CT100_ (paired t-test; n=14). Data are mean ± SEM. *p < 0.05.

**Figure 4.**
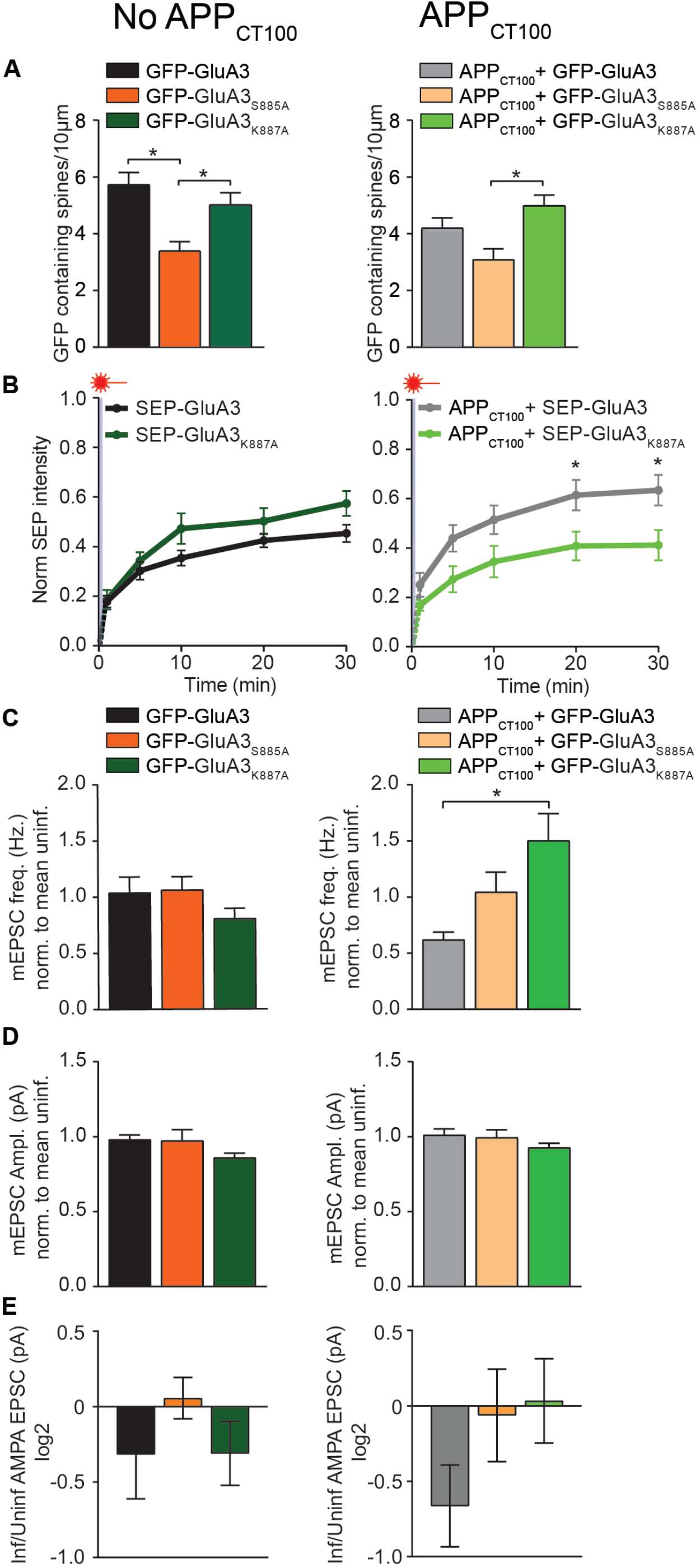
A single amino acid in the PDZ binding motif of GluA3 determines the susceptibility of synapses for Aβ. Comparison of GFP-GluA3 (Fig. 1), GFP-GluA3_S885A_ (Fig. 2) and GFP-GluA3_K887A_ (Fig. 3) under basal and Aβ conditions. (A left) Density of GFP-containing spines was lowered in GFP-GluA3_S885A_ expressing GluA3-KO dendrites compared to those expressing GFP-GluA3 or GFP-GluA3_K887A_ (ANOVA p=0.002; GFP-GluA3 vs GFP-GluA2_S885A_ p=0.001; GFP-GluA3 vs GFP-GluA3_K887A_ =p0.361; GFP-GluA3_S885A_ vs GFP-GluA3_K887A_ p=0.031) (right) During APP_CT100_ expression, GFP-GluA3_S885A_ expressing GluA3-KO dendrites had a lower density of GFP-containing spines compared to those expressing GFP-GluA3_K887A_. (ANOVA p=0.008; GFP-GluA3 vs GFP-GluA2_S885A_ p=0.136; GFP-GluA3 vs GFP-GluA3_K887A_ =p0.306; GFP-GluA3_S885A_ vs GFP-GluA3_K887A_ p=0.006) (B left) FRAP in dendritic GluA3-deficient spines expressing SEP-GluA3 and SEP-GluA3_K887A_ was similar (t20 p=0.154; t30 p=0.102), but (right) during APP_CT100_ co-expression, FRAP of SEP-GluA3 expressing GluA3-deficient neurons was higher (t20 and 30 p= 0.048) compared to those expressing SEP-GluA3_K887A_ (GluA3_K887A_ x APP_CT100_ 2-way ANOVA interaction p<0.01) (C left) Expression of GFP-GluA3, GFP-GluA3_S885A_ and GFP-GluA3_K887A_ similarly affected mEPSC frequency compared to their uninfected neighbors (ANOVA p=0.266), but (right) upon APP_CT100_ co-expression, GFP-GluA3 had a lower mEPSC frequency compared to GFP-GluA3_K887A_ but not GFP-GluA3_S885A_ (ANOVA p=0.006; GFP-GluA3 vs GFP-GluA2_S885A_ p=0.104; GFP-GluA3 vs GFP-GluA3_K887A_ =p0.0.005; GFP-GluA3_S885A_ vs GFP-GluA3_K887A_ p=0.639). (D left) mEPSC amplitude was unaffected by any GFP-GluA3 variant (ANOVA p=0.078), (right) irrespective of APP_CT100_ expression (ANOVA p=0.327). (E left) Changes in the eEPSC amplitude ratio between infected and uninfected GluA3-deficient neurons were not significant in the absence (ANOVA p=0.454) or (right) presence of APP_CT100_ expression (ANOVA p=0.17). Data are mean ± SEM. *p < 0.05.

We here show that the sensitivity of synapses to Aβ depends on the presence of AMPAR subunit GluA3 by expressing GluA3 in CA1 neurons from GluA3-knockout mice. We previously established that virally expressed GFP-GluA3 is present at synapses as GluA2/3-heteromers and not as GluA3-homomers (Renner et al., 2017), which is in agreement with the poor ability of GluA3 subunits to form homomeric receptors (Coleman et al., 2016, Rossmann et al., 2011). Although present at synapses, GluA3 did not lead to an increase in synaptic currents. GluA3-containing AMPARs have a low open-probability and channel conductance compared with GluA1-containing AMPARs and therefore contribute little to synaptic currents (Renner et al., 2017). In addition, GluA2/3s can gradually replace AMPARs at synapses (Shi et al., 2001, McCormack, Stornetta & Zhu, 2006), negating an increase in synaptic strength. Expression of GFP-GluA3 can actually lead to synaptic depression (Shi et al., 2001), possibly as a consequence of low-conductive GluA2/3s replacing high-conductive GluA1/2s at synapses. We found that GluA3 needs to bind GRIP to be stably expressed at dendrites and spines, consistent with the observation that GRIP is required for AMPAR transportation along dendrites and insertion into synapses (Setou et al., 2002, Osten et al., 2000). GluA3 levels were not affected by a mutation in its PDZ-binding motif (GluA3_K887A_) that maintains GRIP binding but disables its phosphorylation (Seidenman et al., 2003, Chung et al., 2000, Kreegipuu et al., 1998), suggesting little phosphorylation of GluA3 by PKCα under basal conditions.

We here demonstrate that an intact PDZ-binding motif in GluA3 is essential for Aβ to trigger synaptic depression. Previous studies showed that Aβ-mediated synapse loss depends on PKCα activity and PICK1 to interact with the PDZ-binding motif of AMPARs (Hsieh et al., 2006, Alfonso, S. I. et al., 2016, Alfonso, S. et al., 2014). We here establish that this applies selectively for the PDZ-binding motif at the c-tail of GluA3. In line with our observations, previous studies observed that PICK1 can mediate the endocytosis and lysosomal targeting of GluA2/3s without affecting GluA1-containing AMPARs (Koszegi, Fiuza & Hanley, 2017, Daw et al., 2000). Possibly PICK1 needs to bind all four AMPAR subunits within an AMPAR-complex to initiate its endocytosis. Alternatively, the GluA1 subunit may inhibit PICK1-mediated AMPAR endocytosis, for instance by recruiting PSD-95 to synapses, which protects them from effects of Aβ (Dore et al., 2021). We hypothesize that Aβ sets a signaling cascade in motion that leads to phosphorylation the GluA3 c-tail, whereupon GRIP detaches from synaptic GluA3, allowing PICK1 to bind and selectively remove GluA2/3s from synapses.

In this study we used organotypic slice cultures from immature mice as a model system, raising reservations about its relevance for AD pathophysiology at advanced age. However, we previously showed that spine loss and memory impairment in aged APP/PS1 transgenic mice require the presence of AMPAR subunit GluA3 (Reinders et al., 2016). Furthermore, GluA3 has been implicated to be associated with AD (Carter et al., 2004, Bodily et al., 2016, Boyle et al., 2006, Thorns et al., 1997, Bereczki et al., 2018). We therefore propose that targeting protein interactions at the PDZ-binding motif of GluA3 may open a potential therapeutic approach against AD.

## Materials and Methods

### Mice

The GluA3-(genetic knock-out) and wild-type littermate colony was established from C57Bl/6 × 129P2-Gria3tm1Dgen/Mmnc mutant ancestors (RRID:MMRRC_030969-UNC) (MMRRC, Davis, CA), which were at least 20 times backcrossed to C57Bl/6 mice. Mice were kept on a 12 hr day-night cycle (light onset 8 or 7am) and had *ad libitum* access to food and water. All experiments were conducted in line with the European guidelines for the care and use of laboratory animals (Council Directive 86/6009/EEC). The experimental protocol was approved by the Animal Experiment Committee of the Royal Netherlands Academy of Arts and Sciences (KNAW).

### Organotypic hippocampal slice preparation and exogenous protein expression

Organotypic hippocampal slices were prepared from P6-8 mice as described previously (Stoppini, Buchs & Muller, 1991) and used at 7–12 days *in vitro* for electrophysiology and 14-21 days *in vitro* for imaging. For the expression of exogenous GFP, APP_CT100_ and GFP- or SEP-tagged rat GluA3 (flip), GluA3_S885A_ and GluA3_K887A_, the respective constructs were cloned into a pSinRep5 shuttle vector. The resulting pSinRep5 plasmids were used to produce infective Sindbis pseudo viruses according to the manufacturer’s protocol (Invitrogen BV). Sindbis virus infection was achieved by injecting diluted virus into slices 20-52 hours prior to the experiments.

### Electrophysiology

During recordings, slices were perfused with artificial cerebrospinal fluid (ACSF): (in mM) 118 NaCl, 2.5 KCl, 26 NaHCO_3_, 1 NaH_2_PO_4_, supplemented with 4 MgCl_2_, 4 CaCl_2_, 20 glucose at 27°C, gassed with 95%O_2_/5%CO_2_. Patch recording pipettes were filled with internal solution containing (in mM): 115 CsMeSO_3_, 20 CsCl, 10 HEPES, 2.5 MgCl_2_, 4 Na_2-_ATP, 0.4 Na-GTP, 10 Na-Phosphocreatine, 0.6 EGTA. Whole-cell recordings in were made with 2.5–4.5 MΩ pipettes (R_access_ < 20 MΩ, and R_input_ > 10× R_access_). During mEPSC recordings TTX (1 μM; Tocris) and picrotoxin (100 μM; Sigma) were added. During evoked recordings, a cut was made between CA1 and CA3, and picrotoxin (50 μM) was added to the bath. Two stimulating electrodes, (two-contact Pt/Ir cluster electrode, Frederick Haer), were placed between 100-200 μm down the apical dendrite and 100-300 μm apart laterally. Two neighbouring Sindbis infected and uninfected CA1 neurons were simultaneously recorded. AMPAR-mediated EPSCs were measured as the peak inward current at ™60 mV directly after stimulation. Data was acquired using a Multiclamp 700B amplifier (Molecular Devices). Mean EPSC amplitudes contained at least 20 sweeps at each holding potential and were acquired using pClamp 10 software (Molecular Devices). mEPSC data are based on at least 100 events or 10 min of recording and analysed with MiniAnalysis (Synaptosoft). Individual events above a 5pA threshold were manually selected by an experimenter blind to the experimental condition.

### Two-photon imaging

3D images were collected by two-photon laser scanning microscopy (Femtonics Ltd.) with a mode-locked Ti:sapphire laser (Chameleon; Coherent) tuned at 910 nm using a 20× objective. During imaging, slices were kept under constant perfusion of ACSF at 30°C, gassed with 95%O_2_/5%CO_2_. For spine densities, apical dendrites were imaged ∼180 μm from the cell body (pixel size x,y,z 0.05 × 0.05 × 0.75μm). The density of spines protruding in the horizontal (*x*/*y*) plane were manually quantified from projections of stacked 3D images by an experimenter blind to experimental condition. For analysis and example images, the intensity value limits of each stacked image were optimized for spine recognition. To monitor the transportation of virally expressed GFP-GluA3 into dendrites, the soma and >150µm of apical dendrite were captured (pixel size x,y,z 0.3 × 0.3 × 0.75μm) using equal microscope settings per condition. For photobleaching experiments, apical dendrites were imaged 150–250μm from the cell body (pixel size x,y,z 0.05 × 0.05 × 0.5μm). Photobleaching of SEP-fluorescence was achieved by prolonged xy scanning of isolated spines for 10-20s, until complete bleaching was visually confirmed (Fig. S3). To determine the fluorescence recovery after photobleaching (FRAP), similarly sized z-stacks of dendrites were collapsed for each time point. Background-subtracted green fluorescence of spines was quantified, normalized to that of its dendrite and compared across time. All image analysis was performed with ImageJ software (fiji.sc).

### HEK-cell co-immunoprecipitation and immunoblotting

Full length GluA3 cDNAs were subcloned into a pRK5-Dest vector, and Grip1 into a pcDNA3.2-V5-Dest vector. HEK cells were passed one day before transfection in DMEM + GlutaMAX (Gibco), 10% FBS (Invitrogen), 1% Penicillin/Streptomycin (Gibco) in 10 cm dishes. 2 h before transfection, the medium of ∼60% confluent cells was refreshed. Cells were transfected with ∼2.5ug Grip-V5 and GluA3, GluA3_S885A_, or GluA3_K887A_ using PEI 2500. The amount of DNA used for transfection with GluA3 constructs was optimized based on protein expression levels beforehand. After ∼48 hours, cells were harvested in 1 ml of a 2% Triton X-100 ice-cold immunoprecipitation buffer (25mM HEPES/NaOH, pH7.4, 150 mM NaCl) containing 2% Triton X-100 and EDTA-free protease inhibitor cocktail (Roche). The resulting samples were incubated for 1 h at 4°C and spun down twice at 20,800 x g for 10 min at 4°C. Anti-Grip (4 µg ABN27, Millipore) was added to the supernatants and incubated overnight at 4°C. The next day, protein A/G PLUS-agarose beads (40 µL; Santa Cruz Biotechnology, Inc.) were added for 1 h at 4°C and washed 4 times with immunoprecipitation buffer containing 1% Triton X-100. Proteins were eluted in SDS sample buffer (55 µL), boiled for 5 min and loaded on a 4-15% Criterion TGX Stain-Free precast gel (BioRad). Protein samples were transferred unto a PVDF membrane (BioRad) overnight at 40V. The blots were blocked in 5% milk in TBST and incubated with primary and secondary antibody in 3% milk in TBST. The following antibodies were used: anti-GluA2/3 (1:2000; CQNFATYKEGYNVYGIESVKI, custom made at Genscript) (Chen et al., 2014), anti-V5 (1:1000; ab27671, Abcam) in combination with goat-anti-rabbit-HRP (DAKO 1:10000) and goat-anti-mouse-HRP (DAKO, 1:10000). Membranes were developed using ECL femto (Thermo Scientific).

### Quantification and statistical analysis

For each experiment, desired sample size was based on similar experiments performed previously. N numbers represent number of neurons with the exception of FRAP data where it represents synapses (max. 3 synapses/neuron). Each experiment was repeated in at least 4 animals per group. No outliers were excluded. In figure 1-3, experimental conditions that are depicted in the same graph were performed in parallel and within the same animals. When necessary, data sets were log-transformed to obtain normal distributions and homogeneity of variance. Experimental conditions were compared using two-tailed Student *t* tests for two conditions (unpaired, unless otherwise indicated) or an ANOVAs with post-hoc Tukey’s multiple comparison test for more than two conditions. FRAP experiments were analysed on multiple timepoints using the Holm-Šídák multiple comparison correction. Where indicated in the figure captions, two-way ANOVAs were used. *P* values below 0.05 were considered statistically significant.

## Acknowledgements

We thank Tessa Lodder and Hans Lodder for expert technical assistance. This work was supported by Alzheimer Nederland, the Dutch Alzheimer association (H.W.K. and N.R.R.) and Brain Foundation Netherlands, de Hersenstichting (H.W.K.).

## Author contributions

NRR and HWK conceived the project. NRR, SvdS, RVK, KJK performed experiments. KWL, ABS and HWK supervised the project. HWK and NRR acquired funding and wrote the manuscript.

---

**Figure S1.**
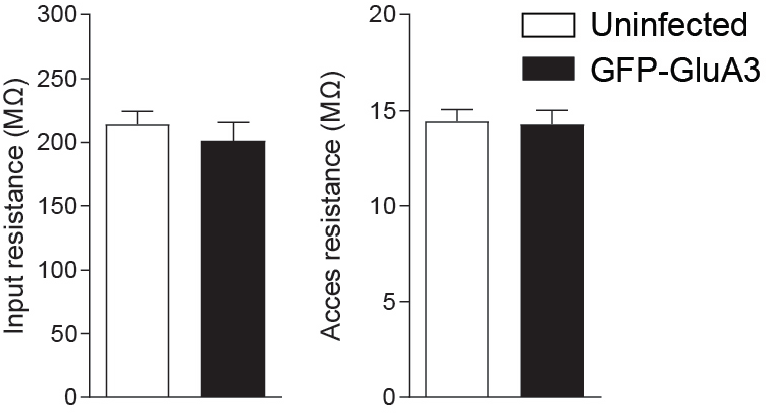
Sindbis infection does not affect basal membrane resistance of CA1 neurons. Dual whole-cell patch clamp recordings of input resistance (left) and access resistance (right) of neighboring GluA3-deficient CA1 neurons either infected with GFP-GluA3 or uninfected, 48 hrs after exposure to Sindbis virus. Data are mean ± SEM.

**Figure S2.**
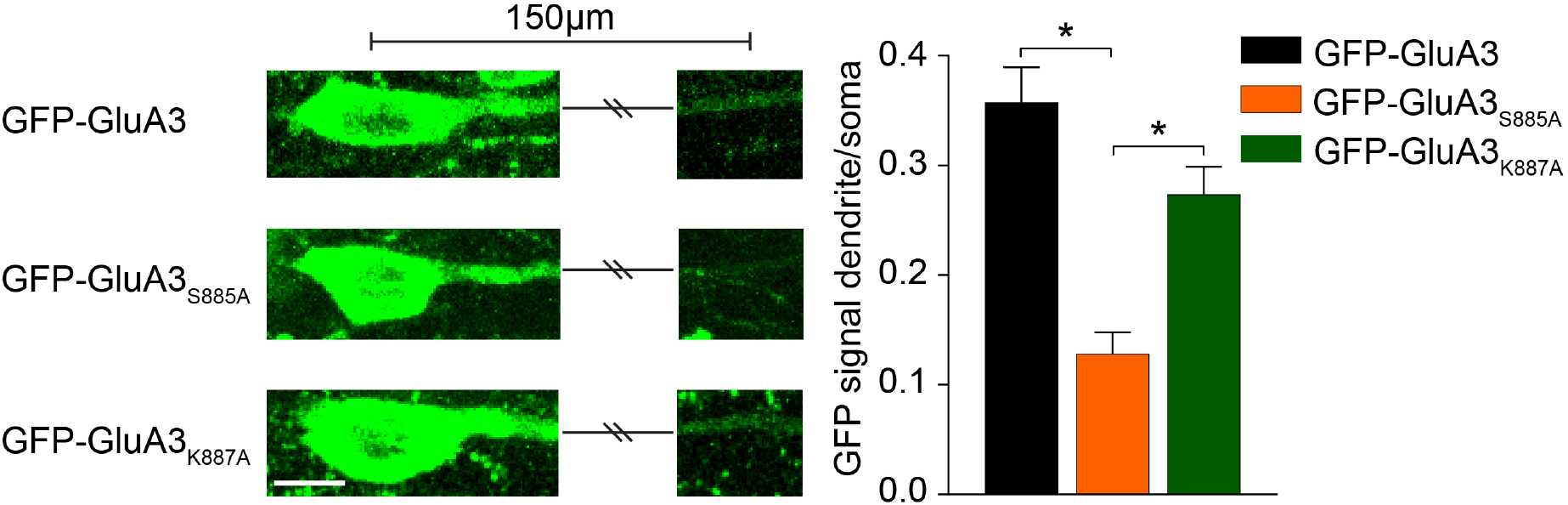
Subcellular distribution of recombinant GFP-GluA3, GFP-GluA3_S885A_ and GFP-GluA3_K887A_ in GluA3-KO CA1 neurons. (left) The GFP intensity in GluA3-KO CA1 neurons expressing GFP-GluA3, GFP-GluA3_S885A_ and GFP-GluA3_K887A_ is low in dendrites compared to soma (images have equal intensity value limits, scale bar= 10µm). (right) The ratio of GFP intensity between dendrite and soma is significantly lower in GFP-GluA3_S885A_ (n=19) expressing neurons compared to those expressing GFP-GluA3 (p<0.001; n=20) or GFP-GluA3_K887A_ (p<0.001; n=20). Data are mean ± SEM. *p < 0.001.

**Figure S3.**
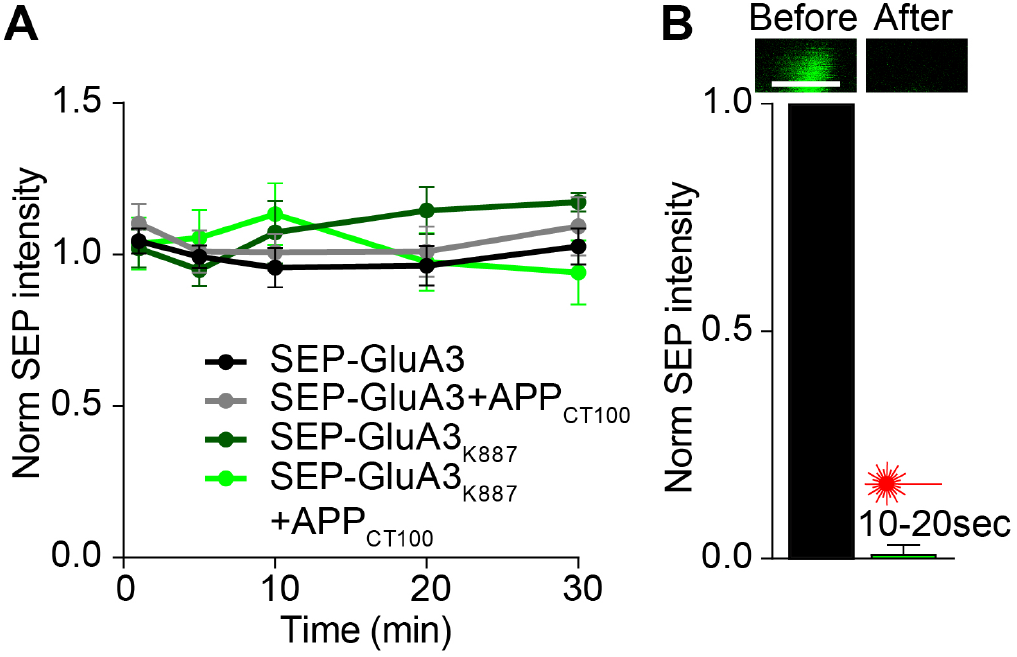
Photo-bleaching of spines minimizes fluorescence intensity without affecting neighboring spines. (A) During FRAP experiments, spines nearby photo-bleached areas showed a stable SEP signal during the experiment (SEP-GluA3, black n=14; SEP-GluA3+APP_CT100_, grey n=16; SEP-GluA3 _K887A_, dark green n=6; SEP-GluA3_K887A_+APP_CT100_, green n=12). (B) SEP signals of spines before and immediately after photo-bleaching demonstrated successful bleaching of SEP fluorescence (n=26 spines, images have equal intensity value limits, scale bar: 1 µm).

**Figure S4.**
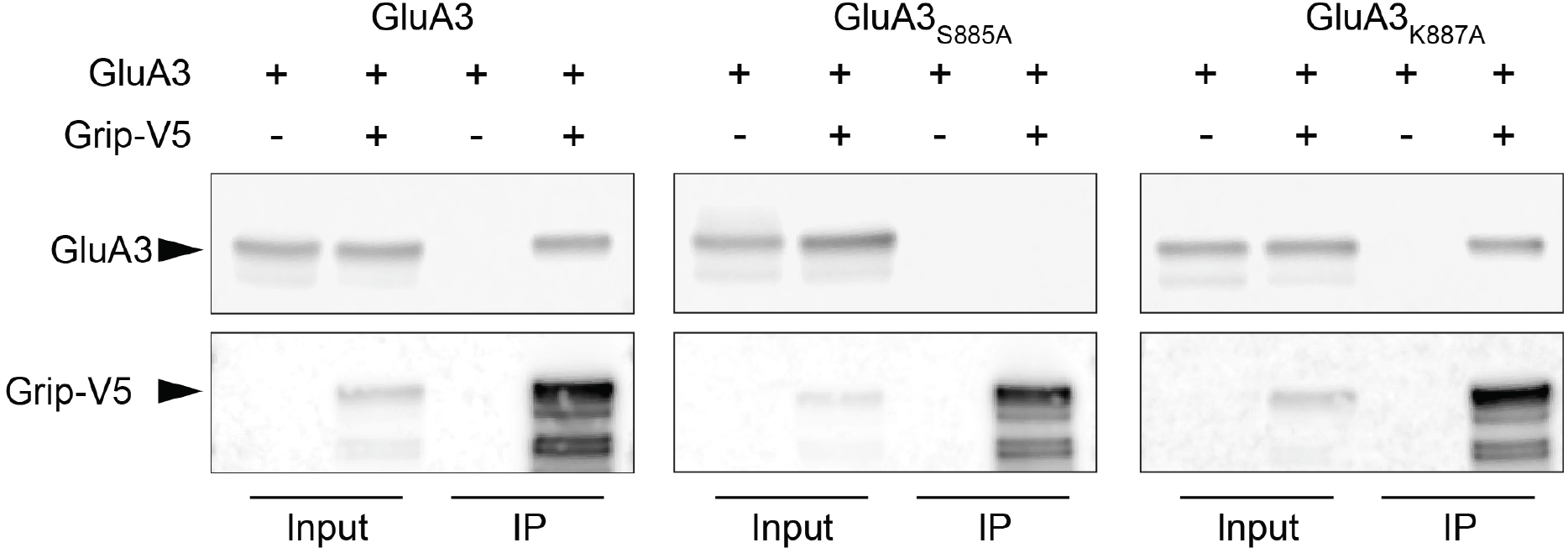
GluA3_S885A_ disables GluA3-GRIP interaction. Western blots from GRIP-V5 IPs on HEK cells expressing a GluA3 variant and/or GRIP-V5. GRIP was immunoprecipitated and stained with an antibody against its fused V5 epitope, GluA3 was stained with anti GluA2/3 antibody. IP = Immunoprecipitation.

## References

Alfonso, S., Kessels, H.W., Banos, C.C., Chan, T.R., Lin, E.T., Kumaravel, G., Scannevin, R.H., Rhodes, K.J., Huganir, R., Guckian, K.M., Dunah, A.W. & Malinow, R. 2014, “Synapto-depressive effects of amyloid beta require PICK1”, The European journal of neuroscience, vol. 39, no. 1460-9568; 0953-816; 7, pp. 1225–1233.

Alfonso, S.I., Callender, J.A., Hooli, B., Antal, C.E., Mullin, K., Sherman, M.A., Lesne, S.E., Leitges, M., Newton, A.C., Tanzi, R.E. & Malinow, R. 2016, “Gain-of-function mutations in protein kinase Calpha (PKCalpha) may promote synaptic defects in Alzheimer’s disease”, Sci.Signal., vol. 9, no. 1937-9145; 1945-0877; 427, pp. ra47.

Bereczki, E., Branca, R.M., Francis, P.T., Pereira, J.B., Baek, J.H., Hortobagyi, T., Winblad, B., Ballard, C., Lehtio, J. & Aarsland, D. 2018, “Synaptic markers of cognitive decline in neurodegenerative diseases: a proteomic approach”, Brain : a journal of neurology, vol. 141, no. 2, pp. 582–595.

Bodily, P.M., Fujimoto, M.S., Page, J.T., Clement, M.J., Ebbert, M.T., Ridge, P.G. & Alzheimer’s Disease Neuroimaging Initiative 2016, “A novel approach for multi-SNP GWAS and its application in Alzheimer’s disease”, BMC bioinformatics, vol. 17 Suppl 7, no. Suppl 7, pp. 268.

Boyle, P.A., Wilson, R.S., Aggarwal, N.T., Tang, Y. & Bennett, D.A. 2006, “Mild cognitive impairment: risk of Alzheimer disease and rate of cognitive decline”, Neurology, vol. 67, no. 1526-632; 0028-3878; 3, pp. 441–445.

Carter, T.L., Rissman, R.A., Mishizen-Eberz, A.J., Wolfe, B.B., Hamilton, R.L., Gandy, S. & Armstrong, D.M. 2004, “Differential preservation of AMPA receptor subunits in the hippocampi of Alzheimer’s disease patients according to Braak stage”, Experimental neurology, vol. 187, no. 2, pp. 299–309.

Chen, N., Pandya, N.J., Koopmans, F.T.W., Castelo-Szekelv, V., van, d.S., Smit, A.B. & Li, K.W. 2014, “Interaction proteomics reveals brain region-specific AMPA receptor complexes”, Journal of Proteome Research, vol. 13, no. 12, pp. 5695–5706.

Chung, H.J., Xia, J., Scannevin, R.H., Zhang, X. & Huganir, R.L. 2000, “Phosphorylation of the AMPA receptor subunit GluR2 differentially regulates its interaction with PDZ domain-containing proteins”, The Journal of neuroscience : the official journal of the Society for Neuroscience, vol. 20, no. 1529-2401; 0270-6474; 19, pp. 7258–7267.

Coleman, S.K., Hou, Y., Willibald, M., Semenov, A., Moykkynen, T. & Keinanen, K. 2016, “Aggregation Limits Surface Expression of Homomeric GluA3 Receptors”, The Journal of biological chemistry, vol. 291, no. 16, pp. 8784–8794.

Daw, M.I., Chittajallu, R., Bortolotto, Z.A., Dev, K.K., Duprat, F., Henley, J.M., Collingridge, G.L. & Isaac, J.T. 2000, “PDZ proteins interacting with C-terminal GluR2/3 are involved in a PKC-dependent regulation of AMPA receptors at hippocampal synapses”, Neuron, vol. 28, no. 0896-6273; 0896-6273; 3, pp. 873–886.

de Wilde, M.C., Overk, C.R., Sijben, J.W. & Masliah, E. 2016, “Meta-analysis of synaptic pathology in Alzheimer’s disease reveals selective molecular vesicular machinery vulnerability”, Alzheimer’s & dementia : the journal of the Alzheimer’s Association, vol. 12, no. 6, pp. 633–644.

Dore, K., Carrico, Z., Alfonso, S., Marino, M., Koymans, K., Kessels, H.W. & Malinow, R. 2021, “PSD-95 protects synapses from β-amyloid”, Cell reports, vol. 35, no. 9, pp. 109194.

Ehlers, M.D., Heine, M., Groc, L., Lee, M.C. & Choquet, D. 2007, “Diffusional trapping of GluR1 AMPA receptors by input-specific synaptic activity”, Neuron, vol. 54, no. 3, pp. 447–460.

Fiuza, M., Rostosky, C.M., Parkinson, G.T., Bygrave, A.M., Halemani, N., Baptista, M., Milosevic, I. & Hanley, J.G. 2017, “PICK1 regulates AMPA receptor endocytosis via direct interactions with AP2 α-appendage and dynamin”, The Journal of cell biology, vol. 216, no. 10, pp. 3323.

Guntupalli, S., Widagdo, J. & Anggono, V. 2016, “Amyloid-β-Induced Dysregulation of AMPA Receptor Trafficking”, Neural plasticity, vol. 2016, pp. 3204519.

Hanley, J.G. 2018, “The Regulation of AMPA Receptor Endocytosis by Dynamic Protein-Protein Interactions”, Frontiers in cellular neuroscience, vol. 12, pp. 362.

Hsieh, H., Boehm, J., Sato, C., Iwatsubo, T., Tomita, T., Sisodia, S. & Malinow, R. 2006, “AMPAR removal underlies Abeta-induced synaptic depression and dendritic spine loss”, Neuron, vol. 52, no. 0896-6273; 0896-6273; 5, pp. 831–843.

Jurado, S. 2018, “AMPA Receptor Trafficking in Natural and Pathological Aging”, Frontiers in molecular neuroscience, vol. 10, pp. 446.

Kessels, H.W., Kopec, C.D., Klein, M.E. & Malinow, R. 2009, “Roles of stargazin and phosphorylation in the control of AMPA receptor subcellular distribution”, Nature neuroscience, vol. 12, no. 1546-1726; 1097-6256; 7, pp. 888–896.

Kessels, H.W., Nabavi, S. & Malinow, R. 2013, “Metabotropic NMDA receptor function is required for beta-amyloid-induced synaptic depression”, Proceedings of the National Academy of Sciences of the United States of America, vol. 110, no. 1091-6490; 0027-8424; 10, pp. 4033–4038.

Kim, C.H., Chung, H.J., Lee, H.K. & Huganir, R.L. 2001, “Interaction of the AMPA receptor subunit GluR2/3 with PDZ domains regulates hippocampal long-term depression”, Proceedings of the National Academy of Sciences of the United States of America, vol. 98, no. 0027-8424; 0027-8424; 20, pp. 11725–11730.

Kopec, C.D., Li, B., Wei, W., Boehm, J. & Malinow, R. 2006, “Glutamate receptor exocytosis and spine enlargement during chemically induced long-term potentiation”, The Journal of neuroscience : the official journal of the Society for Neuroscience, vol. 26, no. 1529-2401; 0270-6474; 7, pp. 2000–2009.

Koszegi, Z., Fiuza, M. & Hanley, J.G. 2017, “Endocytosis and lysosomal degradation of GluA2/3 AMPARs in response to oxygen/glucose deprivation in hippocampal but not cortical neurons”, Sci.Rep., vol. 7, no. 2045-2322; 2045-2322; 1, pp. 12318.

Kreegipuu, A., Blom, N., Brunak, S. & Jarv, J. 1998, “Statistical analysis of protein kinase specificity determinants”, FEBS letters, vol. 430, no. 1-2, pp. 45–50.

Makino, H. & Malinow, R. 2009, “AMPA receptor incorporation into synapses during LTP: the role of lateral movement and exocytosis”, Neuron, vol. 64, no. 1097-4199; 0896-6273; 3, pp. 381–390.

McCormack, S.G., Stornetta, R.L. & Zhu, J.J. 2006, “Synaptic AMPA receptor exchange maintains bidirectional plasticity”, Neuron, vol. 50, no. 0896-6273; 0896-6273; 1, pp. 75–88.

Moretto, E. & Passafaro, M. 2018, “Recent Findings on AMPA Receptor Recycling”, Frontiers in cellular neuroscience, vol. 12, pp. 286.

Osten, P., Khatri, L., Perez, J.L., Kohr, G., Giese, G., Daly, C., Schulz, T.W., Wensky, A., Lee, L.M. & Ziff, E.B. 2000, “Mutagenesis reveals a role for ABP/GRIP binding to GluR2 in synaptic surface accumulation of the AMPA receptor”, Neuron, vol. 27, no. 2, pp. 313–325.

Perez, J.L., Khatri, L., Chang, C., Srivastava, S., Osten, P. & Ziff, E.B. 2001, “PICK1 targets activated protein kinase Calpha to AMPA receptor clusters in spines of hippocampal neurons and reduces surface levels of the AMPA-type glutamate receptor subunit 2”, The Journal of neuroscience : the official journal of the Society for Neuroscience, vol. 21, no. 15, pp. 5417.

Reinders, N.R., Pao, Y., Renner, M.C., da Silva-Matos, C.M., Lodder, T.R., Malinow, R. & Kessels, H.W. 2016, “Amyloid-beta effects on synapses and memory require AMPA receptor subunit GluA3”, Proceedings of the National Academy of Sciences of the United States of America, vol. 113, no. 1091-6490; 0027-8424; 42, pp. E6526–E6534.

Renner, M.C., Albers, E.H., Gutierrez-Castellanos, N., Reinders, N.R., van Huijstee, A.N., Xiong, H., Lodder, T.R. & Kessels, H.W. 2017, “Synaptic plasticity through activation of GluA3-containing AMPA-receptors”, Elife., vol. 6, no. 2050-084; 2050-084.

Rossmann, M., Sukumaran, M., Penn, A.C., Veprintsev, D.B., Babu, M.M. & Greger, I.H. 2011, “Subunit-selective N-terminal domain associations organize the formation of AMPA receptor heteromers”, The EMBO journal, vol. 30, no. 5, pp. 959–971.

Seçil Uyaniker, Sophie J F van Der, Spek Reinders, N.R., Xiong, H., Ka, W.L., Bossers, K., Smit, A.B., Verhaagen, J. & Kessels, H.W. 2019, “The Effects of Sindbis Viral Vectors on Neuronal Function”, Frontiers in Cellular Neuroscience, vol. 13.

Seidenman, K.J., Steinberg, J.P., Huganir, R. & Malinow, R. 2003, “Glutamate receptor subunit 2 Serine 880 phosphorylation modulates synaptic transmission and mediates plasticity in CA1 pyramidal cells”, The Journal of neuroscience : the official journal of the Society for Neuroscience, vol. 23, no. 1529-2401; 0270-6474; 27, pp. 9220–9228.

Selkoe, D.J. 2002, “Alzheimer’s disease is a synaptic failure”, Science (New York, N.Y.), vol. 298, no. 5594, pp. 789–791.

Setou, M., Dae-Hyung Seog, Tanaka, Y., Kanai, Y., Takei, Y., Kawagishi, M. & Hirokawa, N. 2002, “Glutamate-receptor-interacting protein GRIP1 directly steers kinesin to dendrites”, Nature, vol. 417, no. 6884, pp. 83.

Shi, S., Hayashi, Y., Esteban, J.A. & Malinow, R. 2001, “Subunit-specific rules governing AMPA receptor trafficking to synapses in hippocampal pyramidal neurons”, Cell, vol. 105, no. 0092-8674; 0092-8674; 3, pp. 331–343.

Stoppini, L., Buchs, P.A. & Muller, D. 1991, “A simple method for organotypic cultures of nervous tissue”, Journal of neuroscience methods, vol. 37, no. 0165-0270; 0165-0270; 2, pp. 173–182.

Thorns, V., Mallory, M., Hansen, L. & Masliah, E. 1997, “Alterations in glutamate receptor 2/3 subunits and amyloid precursor protein expression during the course of Alzheimer’s disease and Lewy body variant”, Acta Neuropathologica, vol. 94, no. 6, pp. 539–548.

Triller, A. & Choquet, D. 2005, “Surface trafficking of receptors between synaptic and extrasynaptic membranes: and yet they do move!”, Trends in neurosciences, vol. 28, no. 3, pp. 133–139.

Wenthold, R.J., Petralia, R.S., Blahos, J., II & Niedzielski, A.S. 1996, “Evidence for multiple AMPA receptor complexes in hippocampal CA1/CA2 neurons”, The Journal of neuroscience : the official journal of the Society for Neuroscience, vol. 16, no. 0270-6474; 0270-6474; 6, pp. 1982–1989.

